# What are we *really* decoding? Unveiling biases in EEG-based decoding of the spatial focus of auditory attention

**DOI:** 10.1101/2023.07.13.548824

**Authors:** Iustina Rotaru, Simon Geirnaert, Nicolas Heintz, Iris Van de Ryck, Alexander Bertrand, Tom Francart

## Abstract

**Objective:** Spatial auditory attention decoding (Sp-AAD) refers to the task of identifying the direction of the speaker to which a person is attending in a multi-talker setting, based on the listener’s neural recordings, e.g., electroencephalography (EEG). The goal of this study is to thoroughly investigate potential biases when training such Sp-AAD decoders on EEG data, particularly eye-gaze biases and latent trial-dependent confounds, which may result in Sp-AAD models that decode eye-gaze or trial-specific fingerprints rather than spatial auditory attention.

**Approach:** We designed a two-speaker audiovisual Sp-AAD protocol in which the spatial auditory and visual attention were enforced to be either congruent or incongruent, and we recorded EEG data from sixteen participants undergoing several trials recorded at distinct timepoints. We trained a simple linear model for Sp-AAD based on common spatial patterns (CSP) filters in combination with either linear discriminant analysis (LDA) or k-means clustering, and evaluated them both across- and within-trial.

**Main results:** We found that even a simple linear Sp-AAD model is susceptible to overfitting to confounding signal patterns such as eye-gaze and trial fingerprints (e.g., due to feature shifts across trials), resulting in artificially high decoding accuracies. Furthermore, we found that changes in the EEG signal statistics across trials deteriorate the trial generalization of the classifier, even when the latter is retrained on the test trial with an unsupervised algorithm.

**Significance:** Collectively, our findings confirm that there exist subtle biases and confounds that can strongly interfere with the decoding of spatial auditory attention from EEG. It is expected that more complicated non-linear models based on deep neural networks, which are often used for Sp-AAD, are even more vulnerable to such biases. Future work should perform experiments and model evaluations that avoid and/or control for such biases in Sp-AAD tasks.

## 1. Introduction

Auditory attention decoding (AAD) is a well-established term that collectively describes a series of techniques designed to discern which acoustic source a listener is attending to within a mixture of acoustic sources. This is made possible with neural recordings such as electroencephalography (EEG), magnetoencephalography (MEG) or electrocorticography (ECoG) (O’Sullivan et al. 2014, de Cheveigné et al. 2018, Mesgarani & Chang 2012). One of the most sought-after applications of AAD algorithms is their potential to identify the attended speaker in a multi-talker setting, in order to steer the noise suppression algorithms in hearing aids (HAs), which can lead to a novel tier of *neuro-steered HAs*. These devices are currently envisioned to enable effortless control by hearing-impaired users through their brain signals, whereby the attended sounds are automatically recognized and enhanced over non-target sounds, leading to improved speech intelligibility and reduced listening effort expended by the HA users (Geirnaert, Vandecappelle, Alickovic, de Cheveigné, Lalor, Meyer, Miran, Francart & Bertrand 2021, Slaney et al. 2020).

From the rich collection of existing AAD algorithms, two major paradigms stand out: (1) stimulus-reconstruction algorithms which correlate a temporal representation of the acoustic stimulus (e.g., the speech envelope) with neural recordings to select the attended speaker (O’Sullivan et al. 2014, de Cheveigné et al. 2018) and (2) direct classification algorithms which rely solely on neural recordings to distinguish the direction of an attended sound stream (Geirnaert, Francart & Bertrand 2021, Vandecappelle et al. 2021, Su et al. 2022, Pahuja et al. 2023, Cai et al. 2023). The latter is referred to as spatial auditory attention decoding (Sp-AAD), and represents the main focus of this paper. In short, an Sp-AAD decoder is optimized to detect the spatial focus of attention from instantaneous neural features which reflect spatial auditory attention patterns, e.g., lateralization patterns in the spatio-temporal EEG structure. The Sp-AAD decoding paradigm was previously shown to hold two major advantages w.r.t. the traditional AAD paradigms based on stimulus reconstruction: (1) it decodes the direction of an attended sound stream directly from the user’s EEG, i.e., without requiring access to demixed and clean audio signals and (2) it operates accurately on short time-scales (within 1–5 s) (Geirnaert, Francart & Bertrand 2021). These two positive feats make Sp-AAD particularly suitable for real-time attention decoding systems, whereby HA users are exposed to complex acoustic scenes and spontaneously switch their attention between different acoustic targets, often situated at distinct locations.

Yet despite the promising prospects, the precise decoding mechanisms of the Sp-AAD models have not been fully unraveled, nor was their application and generalization validated in a sufficiently realistic Sp-AAD experiment comprising diverse audiovisual scenarios. In particular, several suspicions have been raised regarding potential non-neural biases that could interfere with decoding the direction of auditory attention from EEG (Geirnaert, Francart & Bertrand 2021). In this work, we employed a simple linear method previously introduced for the Sp-AAD task, namely the Common Spatial Pattern (CSP) filtering algorithm (Geirnaert, Francart & Bertrand 2021), in order to investigate two such distinct biases, namely eye-gaze and trial-specific biases, e.g., due to large shifts in the feature space across trials caused by subjects taking a break or ‘recalibrating’ to a new task. Additionally, we evaluated the generalization performance of CSP filters across trials and subjects and incidentally found a third bias that can be leveraged by unsupervised classifiers which aim to cope with across-trial feature shifts. Specifically, we found that even within trials (over a time of less than a few minutes), a substantial drift occurs in the feature space, which could be easily confounded with feature changes caused by changes in spatial auditory attention. Below we further elaborate on the importance and motivations behind studying these biases when it comes to decoding spatial auditory attention.

Firstly, we investigated whether lateralized eye-gaze signals (potentially buried in the EEG) are leveraged by CSP filters, thereby influencing the decoding accuracy. As CSP filters are trained directly on EEG signals during periods of sustained attention to localized speech, it was previously hypothesized they are susceptible to pick up lateralization patterns exhibited by non-neural signals (e.g., eye-gaze, face- or ear-muscle activations) that are correlated to the spatial focus of auditory attention (Geirnaert, Francart & Bertrand 2021, Strauss et al. 2020). Furthermore, a number of recent studies point towards interactions of the oculomotor system and top-down attention-modulated speech processing. For instance, a phenomenon called *ocular speech tracking* was observed by Gehmacher et al. 2023, claiming that gaze activity tracks acoustic features of an attended natural speech signal more strongly than for a distracting sound. The authors also showed evidence that oculomotor activity distinctly contributes to the neural responses of sensors overlapping the auditory processing areas (temporal and parietal). In another study on spatial auditory attention, Popov et al. 2022 showed neurophysiological evidence that alpha power lateralization, an established biomarker for the top-down attention-related mechanism that suppresses distracting input from unattended directions of sound, is closely associated with lateralized oculomotor action, i.e., eye-gaze shifts. However, previous studies employing CSPs in an Sp-AAD task were not able to fully rule out potential contributions from such confounds because these were not explicitly measured or controlled for in the evaluated datasets. Specifically, in the work of Geirnaert, Francart & Bertrand 2021, the AAD dataset on which the CSP filters were evaluated potentially had a strong eye-gaze confound enabled by the presentation of visual cues on the same side with the attended acoustic target (Das et al. 2020). As such, the AAD protocol design in that dataset made it intrinsically difficult to disentangle eye-gaze and attention and draw firm conclusions on whether neural or non-neural mechanisms drive the high CSP decoding accuracies observed in both the subject-dependent and subject-independent analyses. As a follow-up to that work, we here designed a new audiovisual AAD (AV-AAD) experiment to explore whether Sp-AAD is still possible in the case of incongruent eye-gaze. This new protocol comprises experimental conditions where various degrees of correlation were imposed between the spatial directions of to-be-attended *visual* and *auditory* targets, such that they were either co-located, spatially uncorrelated, or the visual stimulus was totally absent.

Secondly, we aimed to explore whether Sp-AAD models are sensitive to overfitting to trial-specific fingerprints (other than eye-gaze) which could potentially stem from non-stationarities in the EEG data, causing the signal statistics to change across trials. This could result in a shift in the feature space when comparing the feature vectors between two EEG segments with a significant amount of time between them. As a result, different trials might populate different locations in the feature space, such that a classifier can recognize a trial based on the location of the extracted features. In a recent review study, Puffay et al. 2023 highlighted and showcased the risk of exploiting such potential trial biases, i.e., trial-specific confounds in the EEG signals that can be recognized by a classifier, allowing it to artificially increase the decoding accuracy (especially in paradigms where a single class label is present per trial). To counteract this, they stress the importance of using a proper cross-validation (CV) scheme such as between-trial data splits (e.g., leave-one-trial-out CV) as opposed to within-trial splits, as the former does not result in artificially high accuracies due to overfitting to latent trial fingerprints. Similar concerns were expressed by Li et al. 2020, who disproved a study that performed an image classification task from EEG signals by finding that a proper train-testing scheme on a block-design EEG dataset degraded the classification accuracies of the trained models to chance, whereas using block-related labels resulted in a high classification accuracy, indicative of the fact that classification pipelines on EEG are susceptible to learn arbitrary block/trial fingerprints occurring in data segments from the same EEG block/trial. Thus, the generalization performance of a model should ideally be probed on data from a held-out trial (Puffay et al. 2023) or subject (Kamrud et al. 2021).

Naturally stemming from the previous point, we lastly evaluated the (in-)ability of the CSP-based Sp-AAD algorithm to generalize across different trials by leveraging the novel AV-AAD dataset, in which multiple EEG trials of the same condition were recorded at two distant points in time. In addition, we investigated the cross-subject generalization performance of the Sp-AAD model, which is especially useful and time-efficient in a practical setting because it would allow the creation of plug-and-play AAD algorithms (i.e., pre-trained on EEG data from previous subjects) that can directly generalize to a new subject with a minimal or even without any calibration session. However, cross-subject generalization could also be sensitive to biases such as eye-gaze, as the EEG patterns related to ocular motion or gaze direction are expected to be fairly similar across subjects. To determine whether eye-gaze signals actually contribute to the generalization performance, our novel AV-AAD paradigm allows the evaluation of generalization across trials and subjects in conditions with and without a confound of eye-gaze. A by-product of this analysis was the finding that within-trial feature drifts significantly degrade the trial generalization performance of the CSP-based Sp-AAD model.

The remainder of this paper is structured as follows: Section 2 describes the experimental setup and the novel AAD audiovisual protocol, Section 3 reviews the CSP-based Sp-AAD algorithm and the hyperparameters used in this study, while Section 4 presents and discusses the obtained results. Finally, conclusions are drawn in Section 5.

## 2. Experimental setup

In this section, we introduce a new dataset that was primarily designed to investigate eye-gaze-related confounds and model generalization in an Sp-AAD task. To this end, different conditions were created, in which the spatial auditory attention is either congruent or incongruent with the eye-gaze.

### 2.1. Participants

Sixteen normal-hearing participants (one male, fifteen females) were recruited to take part in the AV-AAD experiment. The participants’ age ranged between 19-27 years (with mean age and standard deviation of 20.72 *±* 1.00 years). All subjects were native Flemish speakers. Every subject signed an informed consent form approved by the local Medical Ethics committee. Normal hearing for all subjects was verified by pure-tone audiometry.

### 2.2. Audiovisual protocol

For stimulation, we used a self-curated playlist of video clips from”Universiteit van Vlaanderen”‡, a popular platform for science-outreach podcasts delivered by various researchers.

From a database of more than 100 such videos, we pre-selected 30 male-narrated videos with the best audio and video recording quality, spanning a wide variety of scientific topics. To make the experiment as engaging as possible, we let each participant choose their 10 most and 10 least preferred podcasts (with which they were not previously familiar) from our pre-compiled list. To avoid any stimulus pre-exposure effects in the preference selection stage, the subjects were not provided with the actual videos, but only their titles, which summarized the topic. All presented videos were in the .mp4 format and had an original resolution of 1280*×*720 px, but were downscaled to a smaller frame size of 640 x 360 px since smaller versions of the videos will be presented at different locations on the screen (as explained further). The audio tracks of each video were separately extracted, and the overall root-mean-square (RMS) intensity was normalized to -27 dB FS. The silence portions of the audio tracks were not removed nor shortened in order to ensure precise synchronization with the video. Lastly, all used videos and their audio tracks were cut to a duration of 10 min in order to match the trial length.

In total, the experiment included four conditions, each consisting of two trials of 10 min (i.e., each condition lasted for 20 min in total). In each trial, two audio stimuli were simultaneously presented via the RME Fireface UC soundcard (RME, Haimhausen, Germany) connected to two insertphones of type Etymotic ER10 (Etymotic Research, Inc., IL, USA). First, the left and right insertphones were individually calibrated to present sound at an intensity of 65 dB SPL for a frontal source (we used speech-weighted noise as calibration stimulus). To recreate an acoustic spatial impression of sounds coming from distinct left and right locations, each of the two stimuli was separately convolved with head-related impulse responses (HRIRs) corresponding to either −90° or +90° (the HRIRs were measured on a dummy head in an anechoic room§ using in-the-ear microphones, cf. Kayser et al. 2009). This yielded an interaural level difference of approximately 10.6 dB between the ears for a stimulus from +90° degrees.

We randomized the presented stimuli per participant by randomly drawing (without repetition) from the participants’ lists of *most* and *least* preferred podcasts one *to-be-attended* and one *to-be-ignored* speech stimulus, respectively, per experimental trial. For all conditions, the subjects had the same task, i.e., to listen to the target speaker as indicated on the computer screen and to ignore the competing speaker. However, depending on each condition’s type, the subjects were asked to adhere to a different set of visual instructions, explicitly shown at the beginning of each trial.

An overview of the AV-AAD experimental conditions is presented in Table 1. To probe whether eye-gaze has any influence on the CSP decoding of spatial auditory attention in various AV scenarios, these conditions were designed to have different degrees of consistency between the spatial direction of visual and auditory attention. The visual stimulation in each of the four conditions (further referred to by their acronyms) is described in more detail below:

**Table 1:** Overview of the AV-AAD experimental conditions.

| Presentation order | Condition name | Auditory vs. visual attention | Visual task |
| --- | --- | --- | --- |
| 1 | Moving Video (MV) | Incongruent | Follow the *moving video* of the to-be-attended speaker on a randomized horizontal trajectory. |
| 2 | Moving Target Noise (MTN) | Incongruent | Follow the *moving cross-hair* on a randomized horizontal trajectory. |
| 3 | No Visuals (NV) | Incongruent | Fixate on an imaginary point in the center of the *black* screen and minimize eye movements. |
| 4 | Static Video (SV) | Congruent | Fixate the *static video* presented on the same side of the screen as the to-be-attended speaker. |

- In the *Moving Video (MV)* condition, the video of the to-be-attended speaker was presented as randomly moving along a linear horizontal left-right trajectory spanning the entire screen width. The target coordinates of each new video position were randomized along the horizontal axis, and the downscaled video was programmed to move with a constant and moderate speed of 50 px/s between these target points. The random movement was balanced such that the video was presented on each half of the screen for 50% of the time within each trial. Overall, this condition was designed to have complete inconsistency between the spatial visual and auditory attention throughout the whole stimulation duration. Note that despite the lack of spatial correlation, there was still a semantic correlation between the target acoustic and visual stimulus, since the content of the moving video matched the content of the attended speaker.
- The *Moving Target Noise (MTN)* condition visually consisted of a crosshair also randomly moving on a linear horizontal trajectory at 50 px/s, spanning the entire screen width. This condition was intended to be acoustically more challenging than the former, hence background babble noise was added to each insertphone at a Signal-to-Noise Ratio (SNR) of −1 dB (relative to the joint level of both speakers). In addition, by presenting a crosshair instead of a video, the visual semantic cues were removed, in order to enforce the spatial auditory attention as dominant. Altogether, the MTN condition lacked both the spatial and semantic correlation between the audiovisual stimuli.
- In the *No Visuals (NV)* condition, a black screen was presented and the participants were asked to fixate their gaze on an imaginary point within the center of the screen. This was intended as a control condition for visual attention.
- In the *Static Video (SV)* condition, the video of the to-be-attended speaker was statically presented on either the left or the right margin of the screen, in order to match the side of the to-be-attended acoustic stimulus∥. Hence, there was both spatial and semantic correlation between the audiovisual stimuli. This condition was intended as a proxy for most scenarios in daily life, where visual and auditory attention are spatially aligned (i.e., a person directly gazes at the acoustic source they listen to).

In the MV and MTN conditions, the range of the visual angle spanned by the participants’ eyes was geometrically determined from the screen width (47.6 cm) and the average distance from the participant’s head to the screen (45 cm). This yielded an estimation range of (−20°, +20°) relative to the screen center¶.

Time-wise, the conditions were presented in two separate blocks, i.e., the first trial of each condition were presented sequentially, followed by a second block of trials from each condition (cf. fig. 1). As such, the EEG signals belonging to the two trials of each condition were purposefully measured about 40 min apart, in order to capture the EEG non-stationarity across trials, allowing us to probe the between-trial generalization of CSP filters. The conditions were presented in the same order for all subjects (cf. fig. 1). Following preliminary pilot tests, the SV condition was subjectively rated as the least difficult, hence it was deliberately presented last. The MTN condition was not presented to the first three subjects as it was a later addition to the experiment. Hence, the total EEG recording time was 60 minutes (for 3 subjects) and 80 minutes (for the remainder 13 subjects). After every trial of 10 min, there was a short break, which ensured that the participant’s attention levels were constantly refreshed. A longer break of approximately 5 min in-between the two experimental blocks was also offered to the participants. Comprehension questions related to the content of the attended stimulus were presented at the end of each trial, in order to maintain motivation levels and attention throughout the experiment. The participants had to answer with a word or a short phrase, and their behavioral answers were further analysed to monitor task compliance (results are presented in section 4.1).

**Figure 1:**
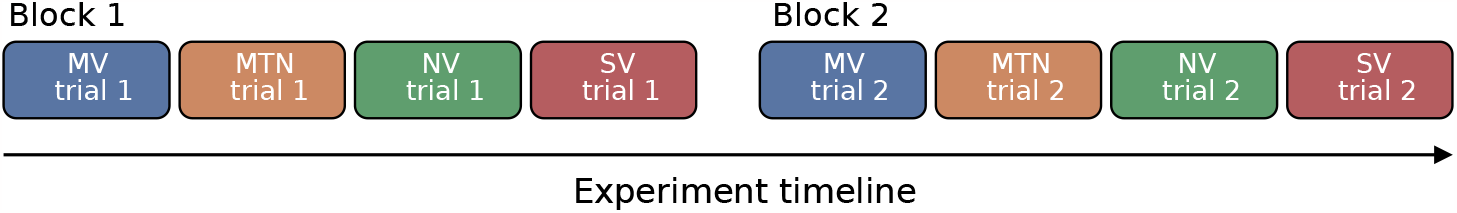
An overview of the experimental timeline in the AV-AAD dataset. A trial is hereby defined as the uninterrupted stimuli presentation for a duration of 10 minutes, during which EEG was recorded. Each color and acronym represents a specific condition, as follows: MV = Moving Video; MTN = Moving Target Noise; NV = No Visuals; SV = Static Video. A block consists of four sequential trials (recorded with small breaks in between), one for each condition type. The trials and conditions were presented in the same order for all participants.

In addition, one *spatial switch of attention* was introduced at the midpoint of each trial (5 min), by programming both stimuli to swap sides from left (L) to right (R) or vice versa. For each subject and condition, the locus of attention in the *first* trial was randomized between (L, R) and (R, L), while the second trial had the opposite attention pair. In conditions with visual stimulation (MV, MTN and SV), an arrow was continuously displayed on the screen, pointing to the direction of the to-be-attended acoustic stimulus. Additionally, the arrowhead changed from L to R (or reversely) after 5 min, to visually cue the participants about the spatial attention switch. Conversely, in the NV condition, the participants were only verbally cued at the beginning of the trial to which speaker (L or R) they had to listen first and were *a priori* instructed to stay alert and switch their attention on the other side when they heard the two stimuli (automatically) swapping sides.

The entire experiment was conducted in a soundproof, electromagnetically shielded room. 64-channel EEG and 4-channel electrooculography (EOG) were recorded at a sampling rate of 8192 Hz with the BioSemi ActiveTwo system (Amsterdam, The Netherlands). The EOG sensors were placed symmetrically around the eyes (cf. fig. 4 in Lopez et al. 2016). Two of the EOG sensors were placed approx. 1.5 cm above and below the right eye (aligned with the center of the eye) to measure vertical oculomotor activity (vertical saccades or blinks). The other two EOG sensors were placed approx. 1 cm right of the right eye and 1 cm left of the left eye, respectively, to measure horizontal oculomotor activity. All visual stimuli were presented on a 21.5 inch screen with a resolution of 1920 *×* 1080 px by running custom-made Python scripts. Synchronization between the audio and video stimuli was performed with the *pygame* module^+^, while synchronization between the measured EEG and the corresponding audio stimuli was achieved via squared-pulse triggers presented every second for the entire duration of each trial and recorded using the EEG system.

## 3. Decoding spatial auditory attention with CSP filtering

### 3.1. CSP filtering

Common spatial patterns (CSP) filtering is a technique widely used in, e.g., BCI motor imagery paradigms to discriminate left-versus right-hand motor imagery (Blankertz et al. 2008, Lotte et al. 2018). Based on previous observations that spatial auditory attention appears to be spatio-temporally encoded in the neural activity (Bednar & Lalor 2018, 2020, Wöstmann et al. 2016, Patel et al. 2018), Geirnaert, Francart & Bertrand 2021 demonstrated the feasibility of CSP filters in an Sp-AAD paradigm to decode the directional focus of auditory attention from EEG in a competing-speaker setting. Below we briefly review the CSP filtering for the binary Sp-AAD paradigm, where the objective is to optimally discriminate between listening to the left and right. For a detailed description of the general theoretical framework of CSP filters, we refer the reader to the studies of Blankertz et al. 2008, Parra et al. 2005.

In a nutshell, the CSP filters **W** ∈ ℝ^*C×K*^ project the (zero-mean) EEG signal **x**(*t*) ∈ ℝ^*C×*1^ measured at a time instance *t* from the original electrode space of *C* channels into a surrogate subspace **y**_*CSP*_ (*t*) = **W**^⊺^**x**(*t*) ∈ ℝ^*K×*1^ of lower dimension *K* ≪ *C*, where the *K* output channels are uncorrelated and the discrimination between the two classes is maximized.

Mathematically, the CSP algorithm determines the orthogonal spatial filters **w**_*k*_ ∈ ℝ^*C×*1^ (the columns of **W**) which maximize the output ratio of variance between the instances of the two classes **x**_1*/*2_(*t*) ∈ ℝ^*C×*1^ in the projected subspace:

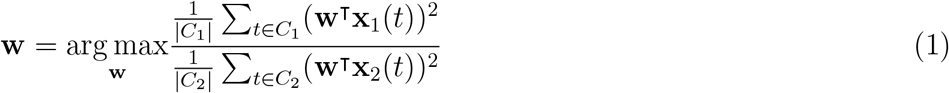

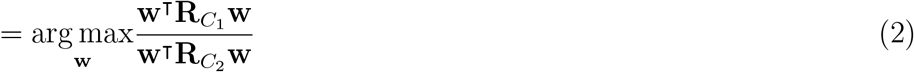

where

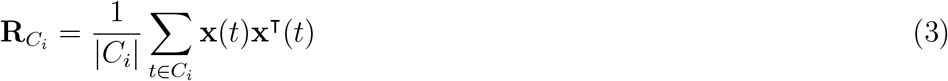

is the covariance matrix of class *C*_*i*_, *i* ∈ *{*1, 2*}* and |*C*_*i*_| denotes the number of time instances in class *C*_*i*_. As shown in Blankertz et al. 2008, the solution of Eq. 2 can be found by computing a generalized eigenvalue decomposition of the class-specific covariance matrices:

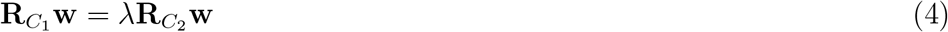

By plugging Eq. 4 into Eq. 2, we obtain:

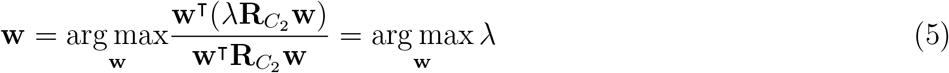

Thus, the best discrimination between the two classes is obtained when the EEG signals are projected onto the generalized eigenvector **w**_1_ corresponding to the largest generalized eigenvalue *λ*_1_. The projection 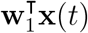 will then produce a virtual channel which will have the maximal relative difference in signal power between the two classes (as targeted in Eq. 1). By reciprocity (i.e., switching the numerator and denominator in Eq. 1), the smallest generalized eigenvector **w**_*K*_ corresponding to the smallest generalized eigenvalue *λ*_*K*_ will achieve the same for the other class. Therefore, to select the *K* most informative spatial filters, the generalized eigenvalues *λ*_*k*_ are sorted and the corresponding first 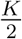 and last 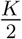 generalized eigenvectors are then selected as columns of **W**.

### 3.2. Classification

CSPs are usually part of a larger classification pipeline to decode directional auditory attention on new EEG data (unseen in the CSP training phase). In the following, we describe the classification procedure which we used in our analysis.

To obtain features for classification, the test EEG data is bandpass-filtered into *B* pre-defined frequency bands and segmented into smaller time windows of length *T*, called *decision windows*. The classification task consists in assigning each decision window to either one of the two classes (attended left or right). To this end, the CSP filters (separately trained per frequency band) are applied per frequency band on each decision window of the test data. Finally, the log-energies of each CSP-filtered window and frequency band are computed (using log-energy features is common in CSP-based classifiers). This results in a total of *B × K* CSP features *f*_*k,b*_ per decision window:

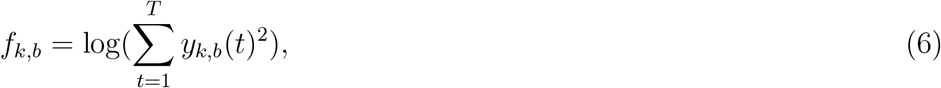

which are stacked together into one feature vector **f** ∈ ℝ^*BK×*1^. To determine the directional focus of attention, **f** is fed to the input of a binary classifier. In this work, we consider two popular classifiers: supervised linear discriminant analysis (LDA) and unsupervised k-means clustering.

#### 3.2.1. Supervised classification with Linear Discriminant Analysis

Fisher’s linear discriminant analysis (LDA) is traditionally used in combination with CSP filters (Lotte et al. 2018). LDA optimizes a linear projection **v** ∈ ℝ^*BK×*1^ within the feature space that maximizes the between-class scatter while minimizing the within-class scatter. This also leads to a generalized eigenvalue problem, which can be solved analytically (Bishop & Nasrabadi 2006):

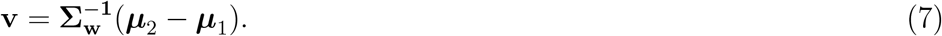

Here, **Σ**_**w**_ ∈ ℝ^*KB×KB*^ is the joint covariance matrix of the features **f** from both classes and ***μ***_1*/*2_ ∈ ℝ^*BK×*1^ are the features’ means across all decision windows in each class. The LDA decision boundary is given by:

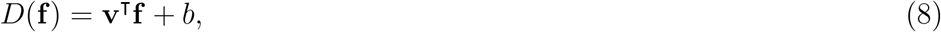

where **v** is defined in Eq. 7, and *b* is the classifier’s bias or threshold, expressed as the mean of the LDA-projected class means:

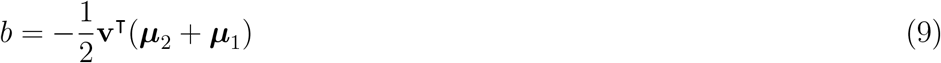

Eventually, **f** is classified into class *C*_1_ if *D*(**f**) *>* 0 and into class *C*_2_ if *D*(**f**) *<* 0.

Note that both CSP and LDA are data-driven and supervised, as they require ground-truth-labeled data to be trained on.

#### 3.2.2. Unsupervised classification with k-means

As noted by Lotte et al. 2018, Huang et al. 2010, using CSPs in combination with LDA classifiers in BCIs could suffer from generalization problems across experimental trials and/or subjects. The potential inability to generalize has been generally attributed to EEG non-stationarities which are unavoidable during long and separate measurement sessions due to changes in the subject’s brain processes, attention and fatigue levels, different artifactual patterns (Blankertz et al. 2007) (e.g., frowning, blinking, swallowing, or yawning) or changes in the experimental conditions (interfering equipment and noise sources).

In an attempt to overcome potential problems with train-test generalization generated by EEG data captured at different time points, we here take a different approach and aim to directly classify the test CSP features in an unsupervised way, thus bypassing any train-data biases in the LDA step. To this end, we replace the supervised LDA with k-means clustering, an *unsupervised* classification algorithm(Bishop & Nasrabadi 2006). In short, k-means finds the optimal partitioning of a given set of observations into K clusters by assigning each observation to the cluster with the nearest centroid (the mean of all points in the cluster) in an iterative process, without needing any *a priori* label/class information.

We note that unsupervised clustering can be performed directly on the test data, i.e., it does not need to be trained on a separate training set (as opposed to CSP and LDA). As a result, it adds flexibility to adapt the classifier to individual (test) trials. However, without further context, it is impossible to then determine which cluster belongs to which class. In Section 3.4, we explain the workaround we employed in computing accuracies for the k-means clustering analyses.

### 3.3. Practical implementation

#### 3.3.1. Data preprocessing

The EEG was initially downsampled using an anti-aliasing filter from 8192 Hz to 256 Hz to decrease the processing time. The EEG trials were then filtered between 1-40 Hz using a zero-phase Chebyshev filter (type II, with 80 dB attenuation at 10% outside the passband) and subsequently re-referenced to the common average of all channels. Afterwards, additional downsampling to 128 Hz was performed to speed up the training of attention decoders. Note that we intentionally refrained from applying z-scoring or any time-related normalization per trial in order to avoid contamination of preprocessing signatures that could be picked up by CSP filters when pooling neighboring data segments across trials, e.g., for cross-validation purposes (see also Section 3.4).

#### 3.3.2. Hyperparameter choices

The CSP filters are usually trained and applied on EEG data filtered in a single frequency band of interest or across a pre-selected range of frequency bands that are relevant to the analysis at hand. In this work, we adopted the so-called filterbank CSP (as in Geirnaert, Francart & Bertrand 2021), which entails that EEG is first filtered into *B* different frequency bands and the CSP filters are then trained and applied per frequency band, resulting in a total of *B* CSP filter matrices **W**_*b*_, with *b* = 1, …, *B*. We considered a filterbank of *B* = 14 overlapping bands spanning a wide frequency range: [1-4], [2-6], …, [24-28], [26-30] Hz. Thus, we intentionally avoided to manually pick a single frequency band, in favor of allowing the classification algorithm to decide which bands are most relevant for distinguishing between left (L) vs right (R) auditory attention.

The number of CSP filters (per frequency band), i.e., *K*, is another tunable hyperparameter. Too many filters are not desirable, as the number of filter weights to optimize increases, and with it, the risk of overfitting. Too few filters are neither optimal, as they might not accurately capture the discrimination between the two classes. Thus, following the conventional CSP recipes from the BCI literature (Blankertz et al. 2008), we chose to train a moderate number of *K* = 6 CSP filters per frequency band.

Before the CSP training step (Eq. 4), we regularized the large-dimensional covariance matrices of each class 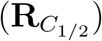 using diagonal loading, i.e., by computing a weighted combination of the sample covariance matrix (potentially poorly-conditioned, but unbiased) and the identity matrix (well-conditioned, but uninformative). The weights of both matrices were automatically determined via the Ledoit-Wolf criterion (Ledoit & Wolf 2004, Geirnaert, Francart & Bertrand 2021).

Finally, as the generalized eigenvalues represent the ratio between class-specific energies of each spatially filtered signal (cf. Eq. 2), they can become corrupted by outlier segments with a high variance. To avoid this issue, the most discriminative CSP filters were selected based on the ratio of median output energies (RMOE) between both classes, cf. Blankertz et al. 2008, (instead of the sorted generalized eigenvalues in Eq. 4), taken over all training windows of length equal to the maximal decision window length that is used in the analysis.

### 3.4. Performance evaluation

The CSP filters were trained and evaluated per experimental condition both within subjects (subject-specifically) as well as across subjects (subject-independently). As evaluation metric, we reported the *decoding accuracy*, i.e., the percentage of correctly classified decision windows averaged across all cross-validation (CV) test folds, CV repetitions and tested subjects. We used several variations of CV schemes in order to discover different biases in the data (see below). In each experiment, the CSP filters and LDA were always trained together on the same subset of data. For different analyses, we used different sets of labels to compute the accuracy (summarized in fig. 2): for probing the eye-gaze bias in Section 4.2, we used labels informative of spatial auditory attention, further denoted as *attention* labels (indicating whether the subject attended to the *L* or *R* speaker); for probing the trial fingerprint bias in Section 4.3, we used labels distinguishing each specific trial, further denoted as *trial* labels (i.e., the EEG segments belonging to the first and second trial of each condition were labelled as 1 and 2, respectively); for probing trial generalization, we used both attention labels and labels informative of feature drift (further details are presented in Section 4.5).

**Figure 2:**
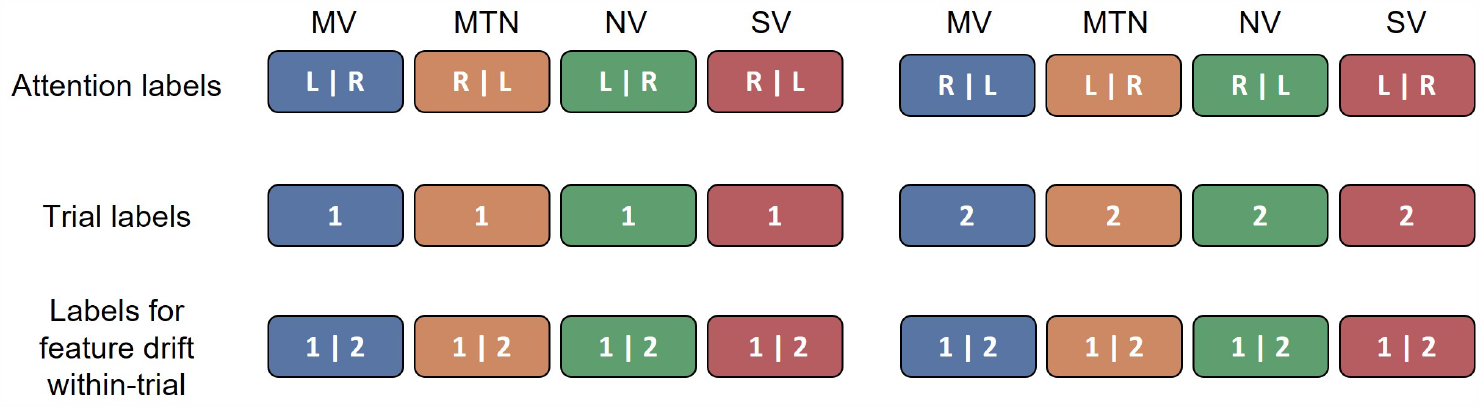
A summary of the different class labels used per condition and trial in different cross-validation settings. The “|” sign delimits the labels assigned to data segments from the first and second half of each trial, respectively. As previously, the colors denote the condition type of each trial.

For the *subject-specific* decoding (further denoted as CSP-SS), separate CSP filters and LDA classifiers were trained per subject and per experimental condition with random 5-fold CV. To this end, the two trials of each condition were concatenated and the resulting EEG data was split into segments of 60 s, which were then randomly shuffled in one of the 5 CV folds, while ensuring a balanced amount of L- and R-labelled segments both in the train and test splits. In order to compute the accuracy obtained with different decision window lengths (WLs), the EEG segments of each test fold were further split up into smaller windows ranging from 1 to 60 s. Per subject, average accuracies were computed across the 5 CV test folds, and across 3 repetitions of the random CV scheme (re-shuffling the 60sec segments into different CV folds). Given that CSP-based classifiers perform comparably well across WLs (Geirnaert, Francart & Bertrand 2021), and that small WLs are required for fast decoding, we used the median accuracies obtained with a WL of 5 s to perform all the statistical analyses presented in Section 4, unless stated otherwise.

We also trained *subject-independent* CSP filters (further denoted as CSP-SI) and LDA classifiers by implementing leave-one-subject-out CV per condition. However, the CSP filters are known to work less well in the subject-independent setting because of the high variability of signals in different frequency bands across subjects (Geirnaert, Francart & Bertrand 2021). Therefore, exclusively for the CSP-SI analysis, the EEG data was bandpass-filtered into one single broadband frequency range (1-30 Hz), and a bias update was applied to the LDA classifier as a normalization step to improve generalization across subjects (details in Geirnaert, Francart & Bertrand 2021).

Furthermore, we investigated CSP generalization across trials by evaluating the CSP-SS decoder with a 2-fold leave-one-*trial* -out CV scheme per condition, leveraging the fact that the two trials of each condition were recorded roughly 40 min apart. Trial generalization was evaluated both with a supervised LDA classifier and with an unsupervised k-means classifier directly applied on the test features in an attempt to compensate for feature drift (in both cases, accuracy was computed based on L/R attention ground-truth labels).

For the classification with k-means clustering, we used *K* = 2 clusters, corresponding to the two attended locations. The cluster centroids were initialized with the (non-deterministic) k-means++ algorithm (Arthur & Vassilvitskii 2006), hence we performed 10 repetitions and up to 1000 iterations to re-update the centroids per repetition. From these 10 repetitions, we selected as final classifier the k-means model with the smallest sums of point-to-centroid distances within-cluster. Notably, k-means is an unsupervised algorithm that assigns arbitrary numerical labels to each cluster (being agnostic to the ground-truth labels), hence it is likely that comparing the clustering output labels to the ground-truth labels results in a low accuracy, which can be attributed to an overall label mismatch. To counteract this, both cluster assignments are tested (i.e., cluster 1 = *L*-attended, cluster 2 = *R*-attended, and vice versa), and the assignment that gives the highest accuracy is retained. Note that this implies we use ground-truth labels, and therefore these results based on k-means classification should not be viewed as representative of a realistic Sp-AAD pipeline. In practice, other heuristics can be used to do this label-to-cluster assignment without the use of the ground-truth labels∗, yet this is beyond the scope of this study, in particular since we will show that even when using this ground truth information, the k-means algorithm is not able to accurately perform Sp-AAD.

For the reported accuracies obtained with LDA classification, the significance level was determined with the inverse binomial distribution, taking into account the total amount of available test data and a significance threshold of *α* = 0.05 (O’Sullivan et al. 2014, Geirnaert, Francart & Bertrand 2021). As the different CV schemes or different WLs have a distinct number of test samples, this results in different significance levels. For k-means clustering, our manipulation of the L/R attended label assignment to each cluster leads to an inherent bias because it “artificially” pulls all the accuracies above 50%, thus also affecting the significance level. To compensate for this bias, exclusively for the k-means classification accuracy, we compute the significance level as the 97.5th percentile of the inverse binomial distribution (which is mathematically equivalent to the 95th percentile of the *folded* inverse binomial distribution, i.e., the true distribution of the biased accuracies - details omitted).

## 4. Results and Discussion

### 4.1. Behavioral results

The subjects’ answers to the comprehension questions were analysed by direct comparison to the correct answers of the attended story in each trial. An average score of 76% correct responses was obtained across all subjects, conditions and trials, indicating a relatively high compliance with the auditory attention task. Per condition, the correct response scores were 75%, 79%, 78% and 71% (across all subjects) for MV, MTN, NV and SV conditions, respectively.

### 4.2. Eye-gaze biases: CSP decoders achieve the highest accuracies in conditions with eye-gaze confounds

The subject-specific CSP (CSP-SS) decoding accuracies following LDA classification with *attention* labels and random 5-fold CV within condition are illustrated in fig. 3a for various WLs. For a WL of 5 s, the obtained median accuracies are 66.4%, 63.8%, 69.6%, 71.4% for the MV, MTN, NV and SV conditions, respectively (fig. 3c). We investigated the effect of condition type on accuracy by means of a Linear Mixed Effects (LME) model where the *condition* was considered as fixed effect and the *subject* as random effect. The LME was fitted to maximize the restricted log-likelihood, and the residuals were checked for normality. Since we were primarily interested in how the audiovisual congruence impacts the decoding accuracy (hence comparing between audiovisual congruent and incongruent conditions), we assigned corresponding contrasts in the model (0.75 for the SV and -0.25 for each of the other conditions). The results revealed that decoding accuracies for the SV condition are significantly higher than for the other conditions (p = 0.002, b = 0.08, CI = [0.03 - 0.13]). This suggests that CSP filters can exploit eye-gaze-related signal components to infer the location of the attended speaker.

**Figure 3:**
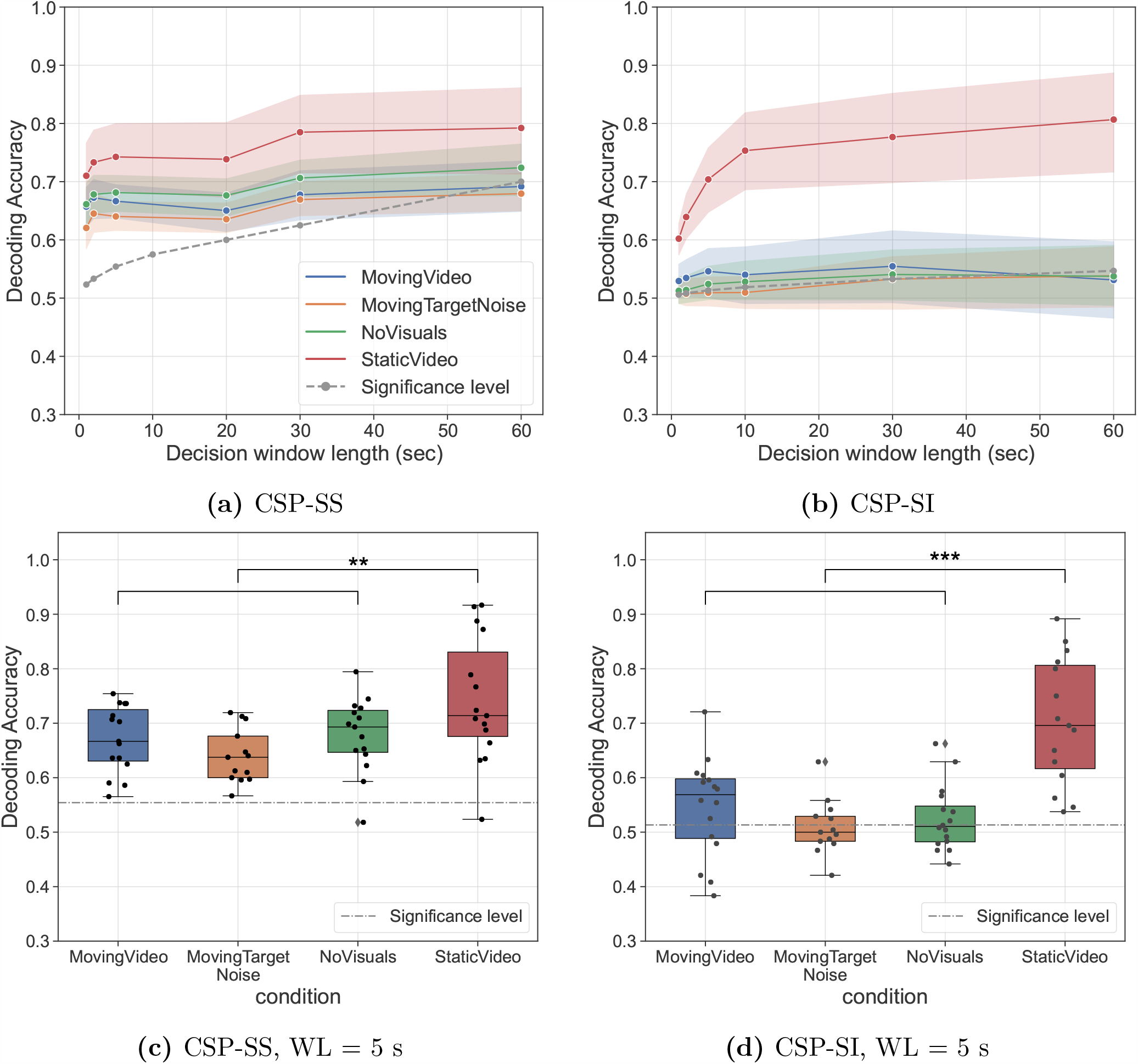
Attention decoding accuracy with subject-specific (CSP-SS) and subject-independent (CSP-SI) decoders peaks for the spatially-congruent audiovisual presentation (SV condition). *Top*: median decoding accuracies for all tested window lengths (WL), with shaded areas representing the 95% confidence interval. *Bottom*: decoding results for WL = 5 s, with each dot representing the accuracy of one subject. Notably, the significance level is reduced for the CSP-SI as more test data was available than for the CSP-SS. The statistical contrast between StaticVideo vs. other conditions’ accuracies was tested with a Linear Mixed Effects (LME) model, whereby (**) marks significant differences for *p* ∈ (1e−3, 1e−2] and (***) for *p* ∈ (1e−4, 1e−3].

**Figure 4:**
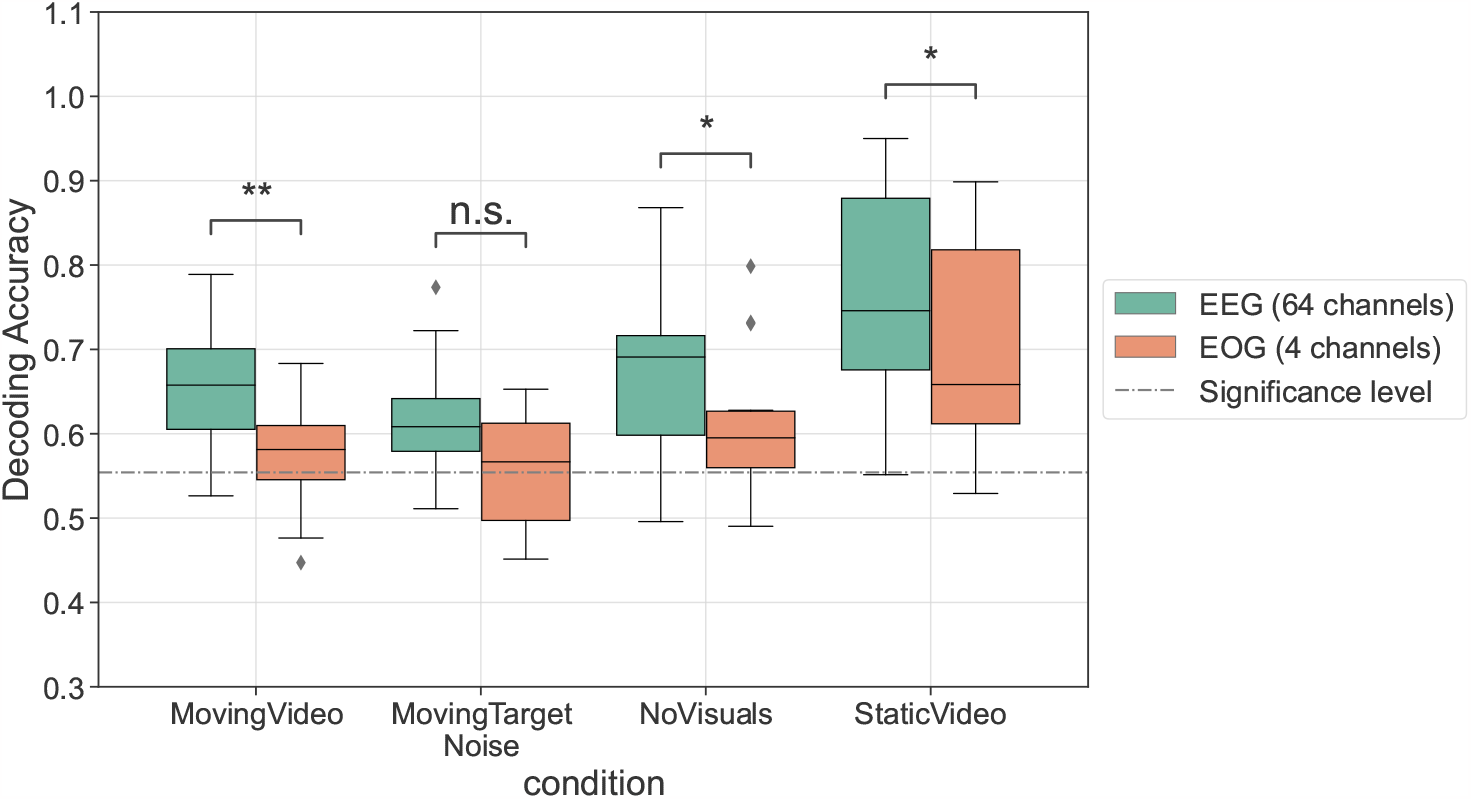
Attention decoding accuracy scores for CSP-SS decoding (with an LDA classifier) on 64 EEG channels vs. 4 EOG channels obtained with random 5-fold CV per condition. To enable a direct comparison between the EEG- and EOG-based CSP decoders, only two CSP filters were trained per frequency band (instead of 6), using attention labels. Statistical significance based on a Wilcoxon signed-rank test is marked with (*) for *p* ∈ (1e−2, 0.05] and with (**) for *p* ∈ (1e−3, 1e−2]. n.s. = not significant.

Additionally, the subject-independent (CSP-SI) decoding results with LDA classification are depicted in fig. 3b. The obtained median accuracies with a leave-one-subject-out CV and a WL of 5 s are 56.9%, 50%, 51.1%, 69.5% for the MV, MTN, NV and SV conditions respectively (fig. 3d). In stark contrast to the CSP-SS decoder, the CSP-SI decoder scores below significance in all audiovisually-incongruent conditions (MV, MTN and NV). In general, poorer performance for a subject-independent model is somewhat expected, as it is more difficult to generalize across subjects than within subject, given the heterogeneous EEG and idiosyncratic CSP feature distributions. Nevertheless, the significant accuracy in the SV condition seems to suggest that CSP is able to capture a dominant subject-independent signal component that is probably related to the eye-gaze direction. An LME model fitted on the CSP-SI accuracies with the same fixed and random effects as for the CSP-SS revealed a similar trend: the accuracy in the audiovisual-congruent SV condition is significantly higher than in the other conditions (*p* ≤ 0.001, b = 0.18, CI = [0.13 - 0.23]). Remarkably, the CSP-SS and CSP-SI accuracies in the SV condition are *not* significantly different (Wilcoxon signed rank test: W=31.5, N=15, p=0.12), possibly suggesting that the eye-gaze signal components are captured similarly well by both the subject-specific and subject-independent decoders. Still, one must interpret this non-significant result with caution, as the underlying amount of training data for the CSP-SI decoders is much higher (300 min, aggregated across subjects) than for the CSP-SS (15 min per train CV fold). As such, it would not be unreasonable to expect that with more training data available, the CSP-SS would score even better than the CSP-SI for the SV condition.

The beneficial effect of the eye-gaze directional information observed in the SV condition could have three possible reasons. Firstly, this result is in line with previous studies, where the spatial alignment of eye-gaze and auditory attention was found to enhance the auditory percept, the behavioral performance in auditory target detection tasks and the discrimination of interaural time and level differences, which are crucial cues for sound localization (Maddox et al. 2014, Pomper & Chait 2017, Best et al. 2007, Andersen et al. 2009). A recent study (Best et al. 2023) showed that gaze direction alone had a strong effect on speech intelligibility: the recall of digit sequences in a competing multi-talker spatial acoustic scene was significantly better when the look direction coincided with the auditory target direction. Moreover, Gehmacher et al. 2023 showed evidence that *ocular speech tracking* (i.e., the phenomenon in which eye-gaze tracks prioritized acoustic features such as the acoustic envelope and acoustic onsets) is more pronounced for an acoustically attended target than for a distractor, thus advocating for a joint network of auditory selective attention and eye movement control. Extrapolating these findings to our Sp-AAD task, it is probable that a visual target spatially aligned with an attended acoustic stimulus enhances the neural auditory attention patterns exploited by CSPs.

Secondly, the AV incongruence, which is artificially enforced in all conditions except in SV, might have made the attention task much harder for the participants to follow. In essence, the incongruent AV conditions are a dual task, with a different visual and auditory spatial focus, which could thus partly explain the significantly lower accuracies observed both for CSP-SS and CSP-SI decoders.

Lastly, it is remarkable that the CSP-SS and CSP-SI accuracies both culminate in the SV condition, and particularly that CSP-SI accuracies are only significant in the SV condition, despite a comparable amount of training data for all conditions. These results thus reinforce the suspicion that CSP decoding is predominantly driven by signal components that originate from the motion of the eyeballs (i.e., EOG-related components), and therefore have no neurological component whatsoever. Assuming this is true, when the look direction does not match with the direction of auditory attention (used as ground truth), the CSP decoding accuracies are expected to drop, which is what we observe in the AV-incongruent conditions. In an additional analysis, we further evaluated this hypothesis by training new CSP-SS filters exclusively on the four external EOG channels (using a similar 5-fold random CV scheme). According to fig. 4, the median decoding accuracies based on EOG channels (with two CSP filters) are consistently lower than those obtained on the standard, 64-channel EEG, regardless of condition. This is confirmed by a non-parametric Wilcoxon signed-rank test (obtained p-values are 0.001, 0.15, 0.03 and 0.04 for MV, MTN, NV and SV respectively). While the EOG-based CSP accuracies are barely significant in the AV-incongruent conditions (as expected, since eye-gaze shifts in those conditions are random), they are above the significance level in the SV condition. This supports the hypothesis that CSP filters do leverage explicit eye-gaze directivity patterns present in both EOG and EEG signals.

While these results make it clear that EOG-related components allow CSP-based decoders to achieve higher accuracies in AV-congruent settings, it remains nevertheless remarkable that decoding accuracies above the significance threshold also occur in conditions without any spatial overlap between the visual and auditory attention (MV, MTN and NV), albeit only in the subject-specific case (fig. 3a, 3c). This suggests that neural patterns purely reflecting the spatial lateralization of *auditory* attention could still be driving the decoding performance in AV-incongruent conditions.

### 4.3. Trial biases: CSP filters can easily discriminate between data coming from distinct trials

fig. 5 illustrates the subject-specific (CSP-SS) decoding accuracies following LDA classification with random 5-fold CV on a WL of 5 s, using *trial* labels (i.e., both the CSP filters and LDA were trained to distinguish between data originating from the two distinct trials of each condition). The median obtained accuracies are 100%, 99.9%, 99.7% and 99.6% for MV, MTN, NV and SV conditions, respectively.

**Figure 5:**
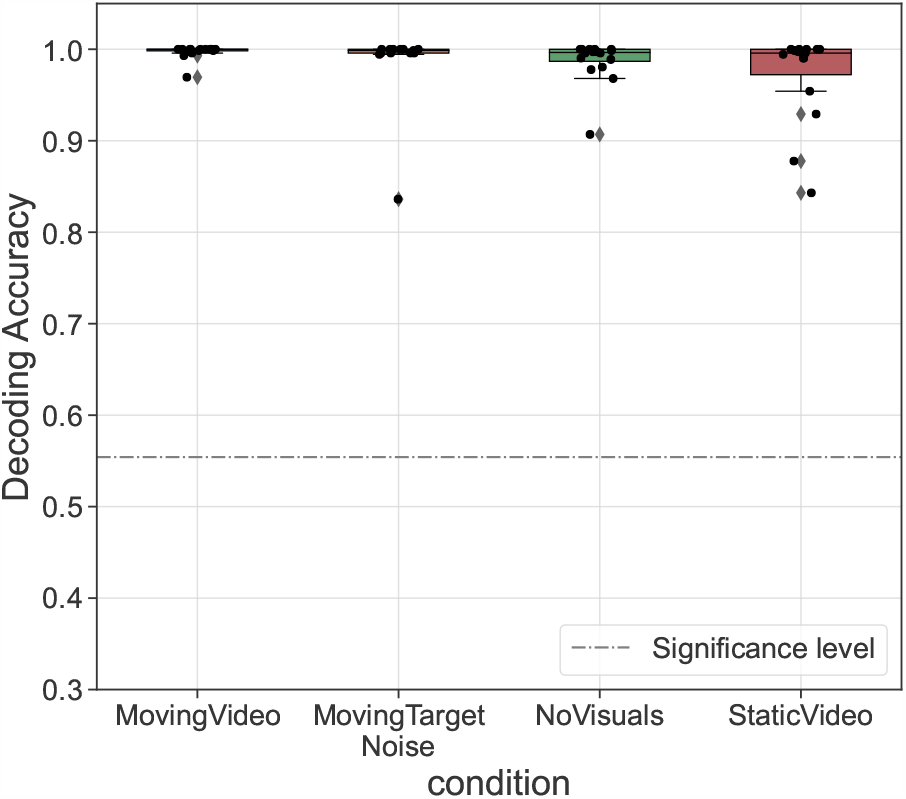
CSP filters in combination with an LDA classifier can discriminate trial fingerprints in all experimental conditions. Depicted are accuracy scores obtained with a CSP-SS decoder and LDA classifier evaluated with random 5-fold CV within condition using *trial* labels (as opposed to *attention* labels) for a WL of 5 s. Each data point corresponds to the CSP-SS accuracy of one subject.

These results confirm the hypothesis that the EEG signals contain trial-specific signatures that can be discriminated by an LDA classifier. The accuracies in fig. 5 are significantly higher than those obtained with attention labels in fig. 3c (Wilcoxon signed rank test: *p* = 3e−5, 2.5e−4, 3e−5 and 6.1e−5 for MV, MTN, NV and SV conditions, respectively), suggesting that trial fingerprints are even more dominant than spatial auditory attention patterns. This implies a strong feature drift over time, since the two trials of each condition were recorded with considerable time in between.

In line with the observations in Puffay et al. 2023, these results mark a red flag for the interpretation of accuracies obtained with random CV, especially when there is a single attention label per trial, which could be easily confounded with the trial label (Su et al. 2022, Pahuja et al. 2023). If the L/R attention labels are balanced within a trial, then one might expect no effect from such biases. However, if there is also a significant feature drift *within* a trial, the classifier can recognize which part of a trial a feature vector comes from, as long as there was training data from a nearby time point.

Note that the benefits from the aforementioned biases are caused by the improper validation based on random CV, in which different short-term windows within a trial are randomly divided between the test and train set. As a result, there is always a high probability that there is one or a few training segments that are close (in time) to the test segment, and therefore have similar time-specific fingerprints that can help to discover the attention label of the test segment. Therefore, in the next subsection, we investigate the Sp-AAD performance in a more correct CV scheme that specifically evaluates across-trial generalization.

### 4.4. Time-related biases: feature shift across trials and feature drift within trials

The results of the CSP trial generalization analysis with supervised LDA and unsupervised k-means are depicted in fig. 6. The median accuracies for the leave-one-trial out (LOTO) evaluation with LDA are largely non-significant (44.6%, 44.26%, 41.45% and 43.3% for MV, MTN, NV and SV conditions respectively, cf. fig. 6a), suggesting that either the CSP or LDA cannot generalize across two separate trials of the same condition and subject. One factor potentially explaining the low accuracies could be the insufficient amount of training data. As each train fold only contains data from one trial (i.e., 10 min), it is rather plausible that a single trial does not display diverse enough EEG signal patterns to enable CSPs to generalize to a similar experimental trial recorded at a later time (despite the fact that the amount of time instances with L/R attention is balanced across trials).

**Figure 6:**
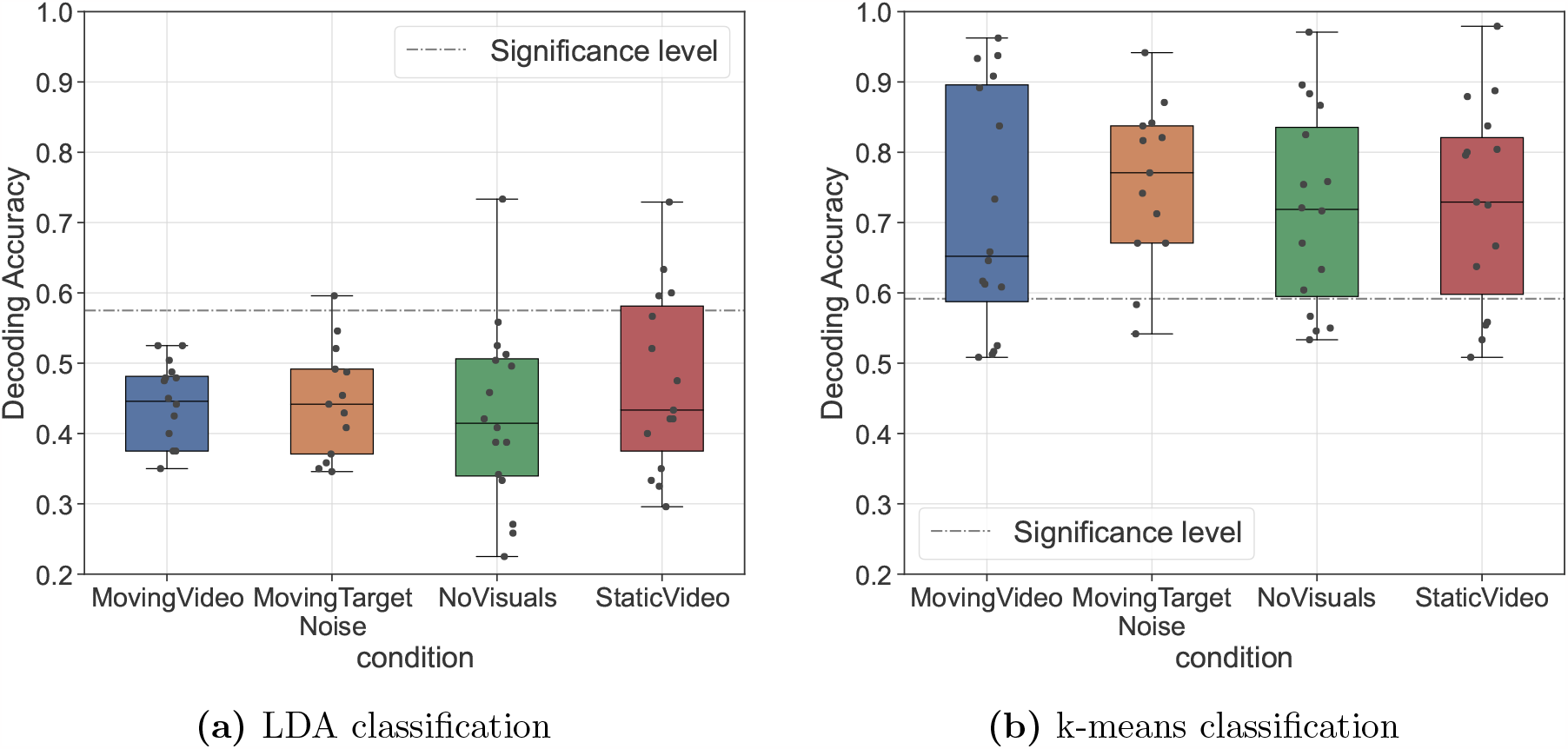
Trial generalization results for decoding the spatial auditory attention with CSP-SS decoders and two different classifiers, i.e., supervised LDA (*left)* and unsupervised k-means (*right*). Each data point represents the accuracy of an individual subject obtained with a *leave-one-trial-out* (LOTO) evaluation per condition (WL = 5 s; *attention* labels are used to train the LDA and to compute the final accuracies for both classifiers).

Nonetheless, the results from fig. 6b confirm that despite the limited amount of train data per condition, the unsupervised k-means classifier is able to restore the median accuracies above the significance threshold, vastly outperforming LDA (the median accuracies obtained with k-means are 65.2%, 77%, 71.9% and 72.9% for MV, MTN, NV and SV conditions, respectively). Thus, it seems that trial generalization with k-means clustering works in all experimental conditions, independently of the presence or absence of the eye-gaze confound. One possible reason is that the k-means algorithm is able to cope with the feature shifts between the training and test trial, as the clustering is performed on the test trial itself. This is further investigated next.

In fig. 7 we present a series of exemplifying CSP feature distributions from the LOTO evaluation, showing a 2D projection of the original high-dimensional CSP feature space following principal component analysis (PCA). It is noteworthy that the trained LDA boundary manages to separate the CSP features of the train trial, but fails to do so for the CSP features of the test trial, as they are shifted or rotated with reference to the features of the train trial. On the other hand, both the train and test feature clouds, if taken separately, still exhibit distinct, well-separable clusters for each class. It thus becomes clear why k-means clustering resulted in the significant accuracies observed in fig. 6b: as k-means is directly applied on the test features, it is completely agnostic to any potential biases from the train data, and can fully leverage the distinct configuration of the test features.

**Figure 7:**
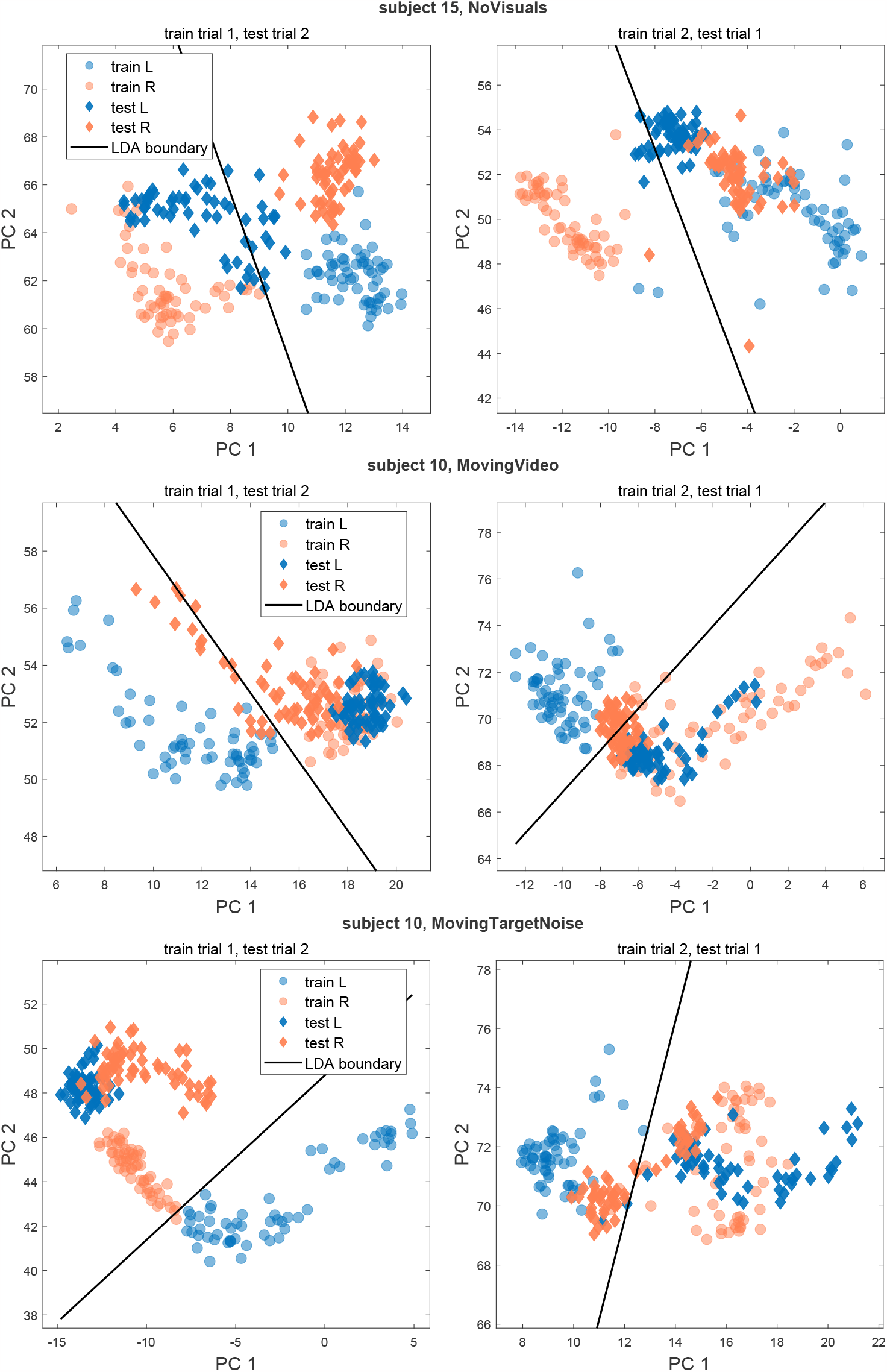
Representative examples of CSP-SS feature shifts and rotations across trials. For 2D visualization, Principal Component Analysis (PCA) was applied on the original CSP features to obtain the first two principal components (PCs) explaining the most variance in the original feature space. Each data point thus corresponds to the first two PCs of every decision window of 5 s. The data points belonging to the train and test sets are marked with circles and diamonds, respectively. The color denotes attention to the left (L) or right (R) speaker. The plotted LDA boundary (solid black line) was obtained after training LDA on the PCs obtained from the CSP train features.

Therefore, if one makes abstraction of the shift between the train and test features, it could be argued that it is possible to create clusters discriminative of spatial auditory attention when trained on one trial from a specific condition and applied to a distinct (test) trial from the same condition, which can be leveraged by an unsupervised clustering algorithm. Yet on a closer look, the CSP features in fig. 7 sometimes appear to drift away in different directions, instead of clustering together in a particular area of the feature space. In fact, depicting the CSP test features as a function of time (fig. 8) reveals that the observed feature drifts are generated by a smooth time-dependent drift in the data. Thus, another potential confound of spatial attention when only considering the features of a single trial are the feature drifts within that trial. With the present dataset, it is unfortunately not possible to conclude if the good k-means accuracies obtained for trial generalization are due to spatial attention (which was alternated only once within any test trial) or due to the time-related feature drifts naturally occurring within that trial.

**Figure 8:**
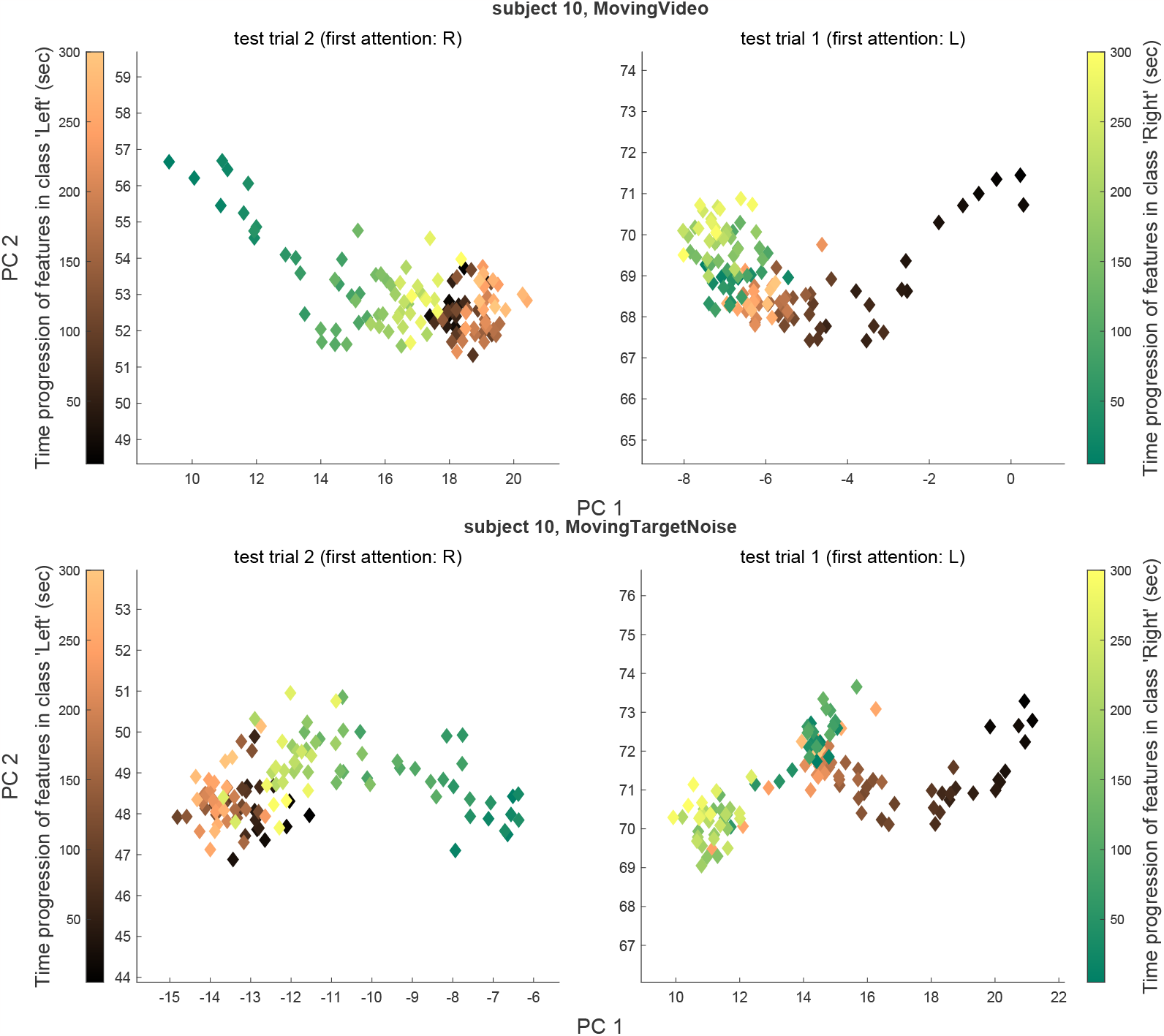
Within-trial feature drifts over time are prevalent in the CSP feature space and they represent a strong confound of the spatial auditory attention in our dataset. The *left* and *right* plots illustrate the 2D-projected CSP features of trial 2 and trial 1, respectively, of two illustrative conditions of subject 10. These features correspond to the *test* CSP features of the latter two conditions plotted in fig. 7. However, they are here color-coded based on their corresponding timestamp within the trial. Thus, features with a lighter color hue (in each attention class) belong to later timestamps, i.e., they occur later in the respective trial. To clarify the relationship between time and attention, we mention as a reference the initial attended location in each trial, and remind the reader that the attended location always changed mid-trial (from *L* to *R* and vice-versa).

To rule out the hypothesis that k-means actually clusters the CSP features’ time-related drifts, one could, for instance, apply k-means on EEG trials where spatial attention is swapped more often than once, and obtain similarly high accuracies as in fig. 6b (when using attention-related labels). Although the trials in our dataset were not designed with frequent switches in spatial attention, this represents a good design feature for future protocols focused on decoding spatial attention. In the following section, we present an alternative analysis meant to probe whether within-trial feature drifts can be classified, thereby introducing yet another potential bias in Sp-AAD studies.

### 4.5. Unsupervised classifiers can leverage within-trial feature drift

To evaluate whether within-trial EEG feature drift can be exploited by unsupervised classifiers, one could in principle apply a classifier on EEG trials recorded during a baseline task, where there is no direct relationship between the time progression and spatial attention, and subsequently compute the accuracy based on labels informative of time progression (e.g., by assigning distinct labels to data segments originating from the *first* and *second half* of each trial, respectively). Since our dataset did not consist of trials that satisfy this condition, we performed this analysis on a separate, publicly-available dataset, namely SparrKULee (Bollens et al. 2023, Accou et al. 2023).

SparrKULee consists of EEG trials recorded from a large sample of 85 normal-hearing subjects, who each performed a basic listening task of attending to continuous single-speaker speech (hence with no alternations in spatial attention). For each participant, a different number of EEG trials with varying lengths were originally recorded. From those, only trials with a duration greater that 10 min were selected and their length was trimmed to exactly 10 min (in order to replicate the experimental conditions in our dataset). These trials were subsequently preprocessed with the same parameters used in our own dataset, as described in Section 3.3.1.

Thereafter, we probed whether k-means can distinguish the feature drift within-trial, namely whether it can separately cluster the CSP features from the first and second half of each test trial. To this end, we applied a similar methodology as in Section 4.4: LOTO cross-validation based on pairs of 10-min trials per subject (i.e., one trial acts as training data and the other as test data). Concretely, we train CSP filters on the *training* trial, and apply k-means on the resulting CSP features in the *test* trial. For both the training and the classification step, we use labels informative of the feature drift within-trial, i.e., labels that distinguish between data segments originating in the first vs. second half of the trial (cf. last row of fig. 2). We note that the CSP training step is rather artificial in this context, as it is only meant to define some arbitrary CSP filters which are applied on the test trial, resulting in test CSP features to be classified by k-means. To obtain pairs of trials which are far away from each other in time (similar to our own dataset), the trials from the first half of the experimental session were paired with the trials from the second half of the experimental session, in chronological order, such that any particular trial occurred only once across all trial pairs.

Fig. 9 reports the average accuracies obtained with k-means clustering over all test trials and across all pairs (per subject), computed in reference to labels informative of within-trial feature drift, for a WL of 5 s. In addition, time-progressive feature drifts similar to those in fig. 8 were also observed in this dataset (figures omitted). The significant accuracies confirm the hypothesis that k-means is able to pick up on feature drifts within-trial that create spurious clusters in the feature space, this being consistent across a large sample of subjects. Extrapolating this interpretation to the previous results on our dataset, it is likely that the significant k-means accuracies obtained in fig. 6b were also driven by the underlying within-trial feature drifts and not by neural patterns related to spatial auditory attention. However, we reiterate that time-related feature drifts and spatial attention cannot be fully disentangled when classifying data from single trials on our dataset, hence the latter claim remains mainly speculative.

**Figure 9:**
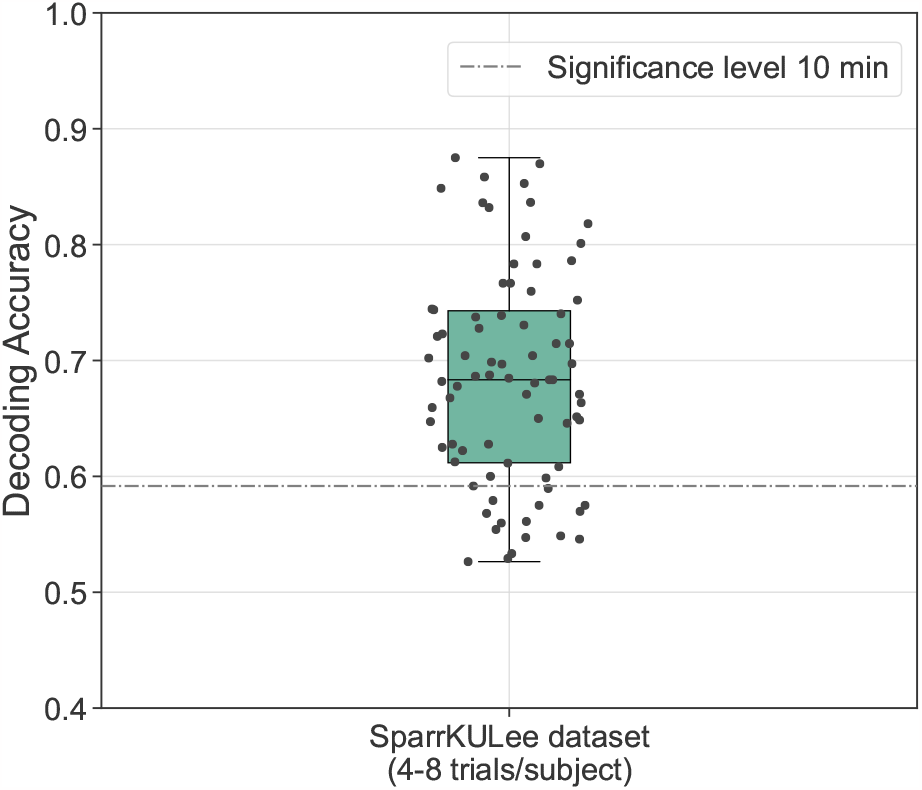
k-means clustering of CSP-SS features can significantly classify feature drifts within-trial on the SparrKULee dataset, which has no confound of spatial auditory attention (the participants underwent multiple trials in which they listened to single-speaker speech). Each dot represents the leave-one-trial-out (LOTO) accuracy of an individual subject for a WL of 5 s, computed based on labels informative of feature drift within-trial.

The main overarching implication of this analysis is that the feature drift within-trial can be decoded and could potentially become a problematic confound when decoding spatial auditory attention across trials with an unsupervised classifier. One way to avoid this confound right from the data collection stage could be through (1) a proper randomization of the order of spatial auditory attention labels across trials and (2) a reduction in the length of segments with sustained attention such that time-related fingerprints are minimally informative for the attention labels.

## 5. Conclusion

In summary, we designed an audiovisual AAD protocol to probe whether spurious signals of non-neural origin interfere with the decoding of spatial auditory attention (Sp-AAD) from EEG. The dataset, comprising EEG recordings from sixteen normal-hearing participants undergoing a spatial auditory attention task, was primarily meant to probe for eye-gaze bias and generalization performance across trials and subjects.

We found that CSP filters trained across- and within-trial are susceptible to capture a whole range of confounding signal patterns. In particular, we showed that lateralized eye-gaze congruent with the spatial target of auditory attention can contaminate EEG signals with such dominant patterns that can be accurately decoded both within and across subjects. Moreover, trial fingerprints and feature drifts within-trial are additional confounds of spatial auditory attention that can be decoded with highly significant accuracies by linear supervised and unsupervised classifiers, such as LDA and k-means clustering. In light of the current results, it still remains an open question whether neural patterns that encode the spatial focus of attention are actually decodable, as the co-occurrence and interplay between these different confounds can profoundly impact even a simple classifier and prevent it from finding EEG signal patterns solely informative of spatial auditory attention.

In order to rule out contributions from such task-irrelevant signals, one would need to design an Sp-AAD protocol where none of these confounds is present, yet this is quite impractical (if not impossible) and will render ecologically invalid and artificial conditions. For instance, spatial alignment between a visual and acoustic target is naturally present in most everyday listening scenarios, while trial fingerprints and entanglement of feature drifts due to either time progression or attention shifts are intrinsic to the measured EEG data. Nonetheless, future work should take measures to mitigate these biases to the best possible extent, such as: inclusion of more frequent switches in spatial attention to ensure a balanced distribution of spatial attention labels per experimental trial; inclusion of explicit EEG preprocessing steps for ocular artifact rejection in order to suppress eye-gaze components from the EEG signals (preferably combined with incongruent audio-visual conditions); proper model evaluation with leave-one-trial/subject-out cross-validation in order to avoid trial-related biases.

While we demonstrated these biases with linear CSP filters and linear classifiers, we believe that more advanced (non-linear) models for decoding spatial auditory attention, such as deep neural networks, can suffer from similar biases, possibly even more due to their higher sensitivity to overfitting (Puffay et al. 2023, Li et al. 2020). Future work developing methodologies for the Sp-AAD task based on EEG should thus be aware of and seek solutions to address these biases before declaring their feasibility and effectiveness.

## 6. Acknowledgements

The authors are grateful to all the participants in this study, and to Anouck Jaspers and Koen van den Eeckhout for their help with data collection. The authors would also like to thank Debora Fieberg for the early brainstorming sessions and for providing the audio stimuli used in this study. Financial support was provided by the Research Foundation Flanders (FWO) (SBO mandate 1S14922N for I. Rotaru, SBO mandate 1S31522N for N. Heintz, SBO mandate 1S34821N for I. Van de Ryck and FWO project G0A4918N), by KU Leuven through a PDM mandate (for S. Geirnaert, No. PDMT1/22/009), and by the European Research Council (ERC) under the European Union’s Horizon 2020 Research and Innovation Programme (Grant Agreements No. 637424 and 802895 for T. Francart and A. Bertrand, respectively). The authors declare no conflicts of interest.

## Footnotes

‡ https://www.universiteitvanvlaanderen.be/

§ We refrained from using HRIRs measured in a non-anechoic room (which include reverberations), because we aimed to make the task perceptually simpler for the participants and allow them to fully focus on the auditory attention lateralization task. This is in line with previous AAD datasets where also anechoic HRIRs were used (Das et al. 2016).

∥ Note there is no 100% correspondence between the direction of the presented video and the direction of attended acoustic stimulus due to physical limitations (e.g., screen size and participant’s distance from the screen): the audio is presented at *±*90° degrees, while the videos are presented at approx. ± 20° relative to the participant’s standpoint, facing the center of the screen (considered at 0 degrees). Thus, both the audio and visual stimuli were presented to either the left or right hemifield of the participants.

¶ A precise measurement was not possible since the participants’ heads were not fixated. As such, they could have spontaneously, even unconsciously, changed their head position during the EEG data collection, thus slightly changing the range of the visual angle.

+ https://www.pygame.org/

∗ e.g., by combining it with a (slower) stimulus reconstruction approach, or based on speaker localization in combination with speaker activity detection.

